# High diversity and sharing of strongylid nematodes in humans and great apes co-habiting unprotected area in Cameroon

**DOI:** 10.1101/2022.09.08.507082

**Authors:** Vladislav Ilík, Jakub Kreisinger, David Modrý, Erich M. Schwarz, Nikki Tagg, Donald Mbohli, Nkombou Irène Charmance, Klára J. Petrželková, Barbora Pafčo

## Abstract

Rapid increases in human populations and environmental changes of past decades have led to intensified contact with wildlife and significantly contributed to pathogen transmission in both directions, especially between humans and non-human primates, whose close phylogenetic relationship facilitates cross-infection. Using high-throughput sequencing, we studied strongylid communities in sympatric western lowland gorillas, central chimpanzees and humans co-occurring in an unprotected area in the northern periphery of the Dja Faunal Reserve, Cameroon. We identified 65 strongylid ITS-2 amplicon sequencing variants (ASVs) in humans and great apes. Great apes exhibited higher strongylid diversity than humans. *Necator* and *Oesophagostomum* were the most prevalent genera, and we commonly observed mixed infections of more than one strongylid species. Human strongylid nematodes were dominated by the human hookworm *N. americanus*, while great apes were mainly infected with *N. gorillae, O. stephanostomum* and trichostrongylids. We were also able to detect rare strongylid taxa (such as *Ancylostoma* and *Ternidens*). We detected eight ASVs shared between humans and great apes (four *N. americanus* variants, two *N. gorillae* variants, one *O. stephanostomum* type I and one *Trichostrongylus* sp. type II variant). Our results show that knowledge of strongylid communities in primates, including humans, is still limited. Sharing the same habitat, especially outside protected areas (where access to the forest is not restricted), can enable mutual exchange of parasites and can even override host phylogeny or conserved patterns.

## INTRODUCTION

Among parasites, strongylid nematodes are of high importance to research as they have infected humans and other vertebrate hosts for millions of years (Anderson, 2000; Dobson & Carper, 1996; Gonçalves, Araújo, & Ferreira, 2003). Strongylid nematodes inhabit various parts of the host body, mainly gastrointestinal and pulmonary tract, where they feed on blood or tissues (Anderson, 2000; Červená et al., 2017; Zajac, 2006). They can live many years within their hosts and generally do not cause mortality; however, severe infections can lead to inflammatory reactions, lesions, severe weight loss, anemia or malnutrition (Delano et al., 2002) and can be attributed to cases of human as well as animal deaths (Lansdale et al., 2017, Roeber et al., 2013). In humans, the most important strongylids are hookworms (*Necator americanus, Ancylostoma duodenale*, and *A. ceylanicum*), infecting over 400 million people worldwide (Loukas et al., 2016). *Necator* hookworms and the nodule worm of the genus *Oesophagostomum* are considered the most prevalent parasites in great apes (Modrý, Pafčo, Petrželková & Hasegawa, 2018).

Unfortunately, identification of distinct strongylid taxa from feces using microscopy is essentially impossible, as strongylid eggs are morphologically indistinguishable (Metzger, 2014). Thus, strongylid identification has been mostly dependent on DNA amplification and sequence analyses (e.g. Cibot et al., 2015; Ota et al., 2015; Santos et al., 2020). Strongylids have mainly been genotyped through DNA amplification methods targeting only one strongylid genus, followed by Sanger sequencing. However, occurrence of complex strongylid communities makes utilization of high-throughput sequencing (HTS) essential (Pafčo, 2017; Pafčo et al., 2018, 2019). HTS of standard phylogenetic markers amplified from complex target populations (metabarcoding) is inexpensive and allows efficient genotyping of hundreds of samples at a time, untangling mixed infections and detecting rare taxa (Valentini, Pompanon, & Taberlet, 2009; Von Bubnoff, 2008; Zhou et al., 2011). Exact delineation of operational taxonomic units (OTUs) can help understand the molecular epidemiology of pathogens and consequently HTS metagenomics has brought about a much deeper insight into the diversity of strongylid nematodes and has revealed hidden zoonotic transmissions or parasite sharing (Avramenko et al., 2017; Lott, Hose, & Power, 2015; Mason et al., 2022; Pafčo et al., 2018).

The close phylogenetic relationship between non-human primates (NHPs) and humans significantly facilitates the overlap and transmission of pathogens and can have a devastating effect on populations of both humans and endangered NHPs (Calvignac-Spencer et al. 2012; Dunay et al., 2018). As the human population is growing rapidly, people are modifying natural animal habitats and contact with wildlife has intensified. Human activities, such as logging and related deforestation, agriculture or “bushmeat” hunting are resulting in a dynamic exchange of pathogens and creating suitable conditions for pathogen transmission (Barlow et al., 2016; Wolfe et al., 2007). Recently, conservation activities and tourism also contribute to transmission of human pathogens to wildlife and can threaten endangered animals (Goldberg et al., 2007; Schwitzer et al. 2019). Several studies have revealed the zoonotic potential of strongylid nematodes with respect to various anthropogenic disturbances; for example, *Oesophagostomum* species were found to be shared between humans and great apes in Eastern Africa (Cibot et al., 2015; Ghai, Chapman, Omeja, Davies, & Goldberg, 2014) and at least two *Necator* species in Central African Republic and Gabon (Hasegawa et al., 2014, 2017). Using the HTS approach, Pafčo et al. (2019) observed hidden transmissions of strongylid nematodes between humans and NHPs in the forest habitats of the Central African Republic, with *Necator* spp. as a main driving force of overlap between different hosts. In this study, we explore strongylid nematode diversity in humans and great apes cohabiting an unprotected area in the northern periphery of the Dja Faunal Reserve (Dja FR), Cameroon. We evaluate possible zoonotic transmission patterns and assess the impact of behavioral/hygiene habits of the local people on their strongylid infections. We employ an ITS-2 metabarcoding approach and predict differences in strongylid nematode communities between different hosts.

## MATERIALS AND METHODS

### Study site, sample collection

Our study took place in the northern periphery of the Dja Faunal Reserve (Dja FR), located in South-East Cameroon. The reserve is a part of the semi-deciduous lowland forest (500–700 m above sea level) with an equatorial and humid climate characterized by one short and one long dry season in between two rainy seasons (February–July/August–November) (Willie, Tagg, Petre, Pereboom, & Lens, 2014). The unprotected area (40 km^2^), comprising the target area of Project Grands Singes (PGS), under Antwerp Zoo Society, Belgium included the research camp La Belgique and three village settlements approximately 25 km from the camp. Several ethnic groups (including the Badjoué, the Fang, the Kaka, the Nzime, the Niem and the Baka) live in the periphery of the reserve in close coexistence with wildlife (Epanda et al., 2020). Although the human population density is low, the pressure on the reserve is substantial, as crops (Arlet & Molleman, 2010), hunting (Ávila et al., 2019; Tagg et al., 2018) and logging (Betti, 2004) remain the main sources of livelihood for the local people. Nevertheless, high densities of central chimpanzees (*Pan troglodytes troglodytes*) and western lowland gorillas (*Gorilla gorilla gorilla*) were recorded in the reserve as well as in the unprotected area around the camp La Belgique (Tagg, Willie, Duarte, Petre, & Fa, 2015).

Human sampling was carried out in three villages – Duomo-Pierre, Malen V, and Mimpala – and great ape samples were collected in secondary forest areas between the villages and in the La Belgique research camp during September and October 2014 (major wet season peak). Fresh fecal samples (total number: *n =* 139) were collected non-invasively from humans (*n =* 48), median age 26 years, and free-ranging great apes: central chimpanzees (*n* = 31) and western lowland gorillas (*n* = 60). Human participants were provided with sampling tubes and samples were then gathered by researchers in the villages. Samples of great apes were collected from the ground under morning nests, with a maximum of three hours after individuals left their nests. To prevent re-sampling of the same individuals and groups of individuals, only groups of different sizes (at the same locality) or groups of the same size (but not coexisting at the same locality) were sampled, and one sample per nest was taken. The samples were immediately fixed in 96% ethanol and stored at room temperature for a maximum of two weeks until they were sent to the Department of Pathology and Parasitology of Veterinary University Brno, where they were stored at −20°C.

Human participants also filled out a questionnaire about their lifestyle including frequency of entering the forest, interaction with great apes, clothing, hygiene, anthelmintic treatment and dietary habits **(Table 3)**. All participants spoke French and researchers assisted them to fill in the questionnaires. Human sampling and data collection followed the protocol approved by the Ethics Committee of the Biological Centre of Academy of Sciences, České Budějovice, Czech Republic and was approved by the local authorities. Sampling was performed after obtaining informed consent of all registered volunteers.

### DNA isolation, library preparation, sequencing

We extracted total genomic DNA from fecal samples using PowerSoil DNA isolation kit (MO BIO Laboratories, Qiagen company, USA) and amplified ribosomal DNA (rDNA), specifically the variable section of rDNA (internal transcribed spacer 2; ITS-2). We prepared sequencing libraries according to the protocol of Pafčo et al. (2018), using two-step PCR following the Fluidigm Access Array primer design. We processed each sample in duplicate, and included two negative and three positive controls according to the protocol. We sequenced the final libraries using the Illumina MiSeq platform (Illumina MiSeq Reagent Kit v2, sequencing 500 cycles of 2 × 250 bp paired-end reads). Additionally, we created a large metadata table containing sample identification (ID), collection site and host species.

### Data processing and statistics

We trimmed raw .fastq sequences using Skewer (Jiang, Lei, Ding, & Zhu, 2014) and followed by paired-end reads assembly in PEAR merger (Zhang, Kobert, Flouri, & Stamatakis, 2014). We eliminated low quality sequences (with expected error rate > 1 %) from the dataset. We detected ITS-2 amplicon sequencing variants (ASVs) and estimated sample relative abundances using software dada2 (Callahan et al., 2016). Using dada2’s algorithm for chimera detection, sequences inconsistently present in both duplicates were marked as chimeras and we removed them from downstream analyses (5–7% of sequences after quality control). We searched for corresponding sequences via standalone BlastN (performed on the NCBI nt database, which was downloaded on 10^th^ February 2020); we excluded environmental or uncultured samples from the database and filtered out all blast hits with < 85% identity and < 90% coverage from the file. We downloaded taxonomy for blast hits using taxize package (Chamberlain & Szöcs, 2013), and assigned taxonomy via dad2, implementing a Naïve Bayesian Classifier algorithm (Q. Wang, Garrity, Tiedje, & Cole, 2007). We merged the resulting taxonomy table with our metadata table into a single phyloseq object, suitable for downstream analyses.

We executed all data analyses in the statistical software RStudio (https://www.rstudio.com). We de-noised the raw dataset (variants unclassified up to “family” level and “non-strongylid” were removed from the dataset) and used a generalized linear model (GLM) with quasipoisson error distribution to test differences in alpha diversity, evaluated as number of ASVs per sample, among the studied hosts. Additionally, we employed post-hoc testing (Tukey) to identify levels of factorial response that differ from each other. Moreover, we measured the alpha diversity by Shannon’s and Simpson’s indexes; we defined community composition as prevalence and relative representation of ITS-2 ASVs using Jaccard and Bray-Curtis ecological distances. In order to prevent negative eigenvalues during computation, we performed square root transformation of the dataset. We then performed Principal coordinate analysis (PCoA) on both Jaccard and Bray-Curtis dissimilarities. To test the interspecific differences in strongylid nematode community compositions among the hosts, we executed permutational analysis of variance (PERMANOVA), followed by analysis of similarity (ANOSIM). We implemented Multivariate general linear models (GLMs) from the R package mvabund (Wang, Naumann, Wright, & Warton, 2012) to search for community-wide divergence and identification of significant ASVs that varied due to the different host species effect. For better resolution, we constructed a diagram showing proportion of reads for significant variants. We further implemented GLM testing with quasipoisson error distribution, followed by PERMANOVA and ANOSIM to evaluate the impact of all factors from the questionnaires **(Table 3)** on the strongylid alpha and beta diversity in humans.

## RESULTS

We analysed fecal samples of humans (*n =* 48), western lowland gorillas (*n =* 60) and central chimpanzees (*n* = 31). In total, 2,943,087 high-quality reads were identified, with a median sequencing depth per sample of 15,612 (minimum = 9, maximum = 375,905). Taxonomic assignment revealed 65 ITS-2 amplicon sequencing variants (ASVs), including at least five strongylid genera **(Table 1)**. Thirty-two unassigned variants (present in 45% of samples) were tentatively classified as being closest to *Nematodirus* sp. or *Travassostrongylus* sp.; however, the sequence identity as well as match scores were low (84.1 % and 76.8 %, respectively) and thus those variants probably do not represent these genera. The last mentioned are among the strongylid nematodes whose sequences are not represented in the database.

**Table 1:** List of identified strongylid nematodes found in studied hosts, sequences NCBI Accession numbers and reference.

| Family | Genus | Species | NCBI Accession | Reference |
| --- | --- | --- | --- | --- |
| **Chabertiidae** | *Oesophagostomum* | *Oesophagostomum stephanostomum* type I | KR149648.1 | Cibot *et al.*, 2015 |
|  | *Oesophagostomum* | *Oesophagostomum stephanostomum* type II | AB821022.1 | Makouloutou *et al.*, 2014 |
|  | *Oesophagostomum* | *Oesophagostomum* sp. | KR149658.1 | Cibot *et al.*, 2015 |
|  | *Ternidens* | *Ternidens deminutus* | AJ888729.1 | Schindler *et al.*, 2005 |
| **Ancylostomatidae** | *Necator* | *Necator americanus* | LC088287.1,<br>LC036563.1<br>MG256601.1 | Hasegawa, 2015; Hasegawa *et al.*, 2017; Jariyapong & Punsawad, 2017 |
|  | *Necator* | *Necator gorillae* | LC088299.1 | Hasegawa *et al.*, 2017 |
|  | *Necator* | *Necator* sp. | AB793535.1 | Hasegawa *et al.*, 2014 |
|  | *Ancylostoma* | *Ancylostoma* sp.<sup>†</sup> | LC036567.1 | Hasegawa, 2015 |
| **Trichostrongylidae** | *Trichostrongylus* | *Trichostrongylus* sp. type I | Unassigned<sup>‡</sup> | NA |
|  | *Trichostrongylus* | *Trichostrongylus* sp. type II | LC185220.1 | McLennan *et al.*, 2017 |
| **Unclassified** | Unclassified | Unclassified | Unassigned<sup>§</sup> | NA |

<sup>†</sup>Closest hit *A. ceylanicum* (similarity 95.5 %)
<sup>‡</sup>Closest hits *T. vitrinus* (similarity 98.48 %), and *T. colubriformis* (similarity 97.34 %)
<sup>§</sup>Probably two taxa: closest hits *Nematodirus* sp. (similarity 84.1 %) and *Travassostrongylus* sp. (similarity 76.8 %)

The most prevalent variants belonged to three genera: *Necator, Oesophagostomum* and *Trichostrongylus* **(Table 2)**. A bar graph visualizing relative abundances of strongylid variants for all studied individuals revealed interspecific differences in the composition of strongylid nematode communities according to host species **(Figure 1)**. Humans were predominantly infected by *N. americanus* (66.7 %), while *N. gorillae* variants were less common (16.7 %). A significant portion of human infections also included *O. stephanostomum* type I (27.1 %). *Trichostrongylus* sp. type II (2.1 %) and four unassigned variants (8.3 %) were also found in humans. Strongylids in great apes were dominated by variants of *N. gorillae* (overall prevalence 91.9%), *Oesophagostomum stephanostomum* type I (89.3%), *Trichostrongylus* sp. type II (69.0%) and unassigned variants (67.9%). *Necator americanus* variants were found only in gorillas (31.7%), while there was no evidence for *N. americanus* in chimpanzees. Additionally, unidentified variants of *Necator* species (neither *N. americanus* nor *N. gorillae*) were detected in great apes (13.3% in gorillas; 9.7% in chimpanzees). Four taxa were recorded in low prevalence solely in gorillas (*Oesophagostomum* sp., *Trichostrongylus* type I, and *T. deminutus*) and one taxon was detected only in a chimpanzee (*Ancylostoma* sp.). We found eight ASVs shared between humans and great apes (8.25% of all observed ASVs), suggesting zoonotic transmission: two *N. gorillae* variants, one *O. stephanostomum* type I, and one *Trichostrongylus* sp. type II variants were found in humans, gorilla and chimpanzees, while four *N. americanus* variants were shared only between humans and gorillas.

**Table 2:** List of numbers of identified amplicone sequence variants (ASVs), their proportion of total reads, numbers of infected hosts and ASVs prevalence among host species.

| Parasite taxa | Number of identified ASVs | Total reads proportion (%) | Number of ASVs in humans | Number of ASVs in gorillas | Number of ASVs in chimpanzees | Prevalence in humans (%) | Prevalence in gorillas (%) | Prevalence in chimpanzees (%) |
| --- | --- | --- | --- | --- | --- | --- | --- | --- |
| *Oesophagostomum stephanostomum* type I | 16 | 41.1 | 1 | 12 | 11 | 27.1 | 81.7 | 96.8 |
| *Oesophagostomum stephanostomum* type II | 3 | 0.7 | 0 | 2 | 3 | 0 | 6.5 | 35.5 |
| *Oesophagostomum* sp. | 1 | > 0.1 | 0 | 1 | 0 | 0 | 1.7 | 0 |
| *Necator americanus* | 16 | 21.7 | 15 | 5 | 0 | 66.7 | 31.7 | 0 |
| *Necator gorillae* | 14 | 20.0 | 2 | 14 | 4 | 16.7 | 96.7 | 87.1 |
| *Necator* sp. | 8 | 0.1 | 0 | 6 | 3 | 0 | 13.3 | 9.7 |
| *Trichostrongylus* sp. type I | 3 | 0.2 | 0 | 3 | 0 | 0 | 13.3 | 0 |
| *Trichostrongylus sp.*<br>type II | 2 | 6.3 | 1 | 2 | 1 | 2.1 | 76.7 | 61.3 |
| *Ancylostoma sp.* | 1 | > 0.1 | 0 | 0 | 1 | 0 | 0 | 3.2 |
| *Ternidens deminutus* | 1 | > 0.1 | 0 | 1 | 0 | 0 | 1.7 | 0 |
| Unclassified | 32 | 10.0 | 4 | 18 | 17 | 8.3 | 58.3 | 77.4 |

**Figure 1:**
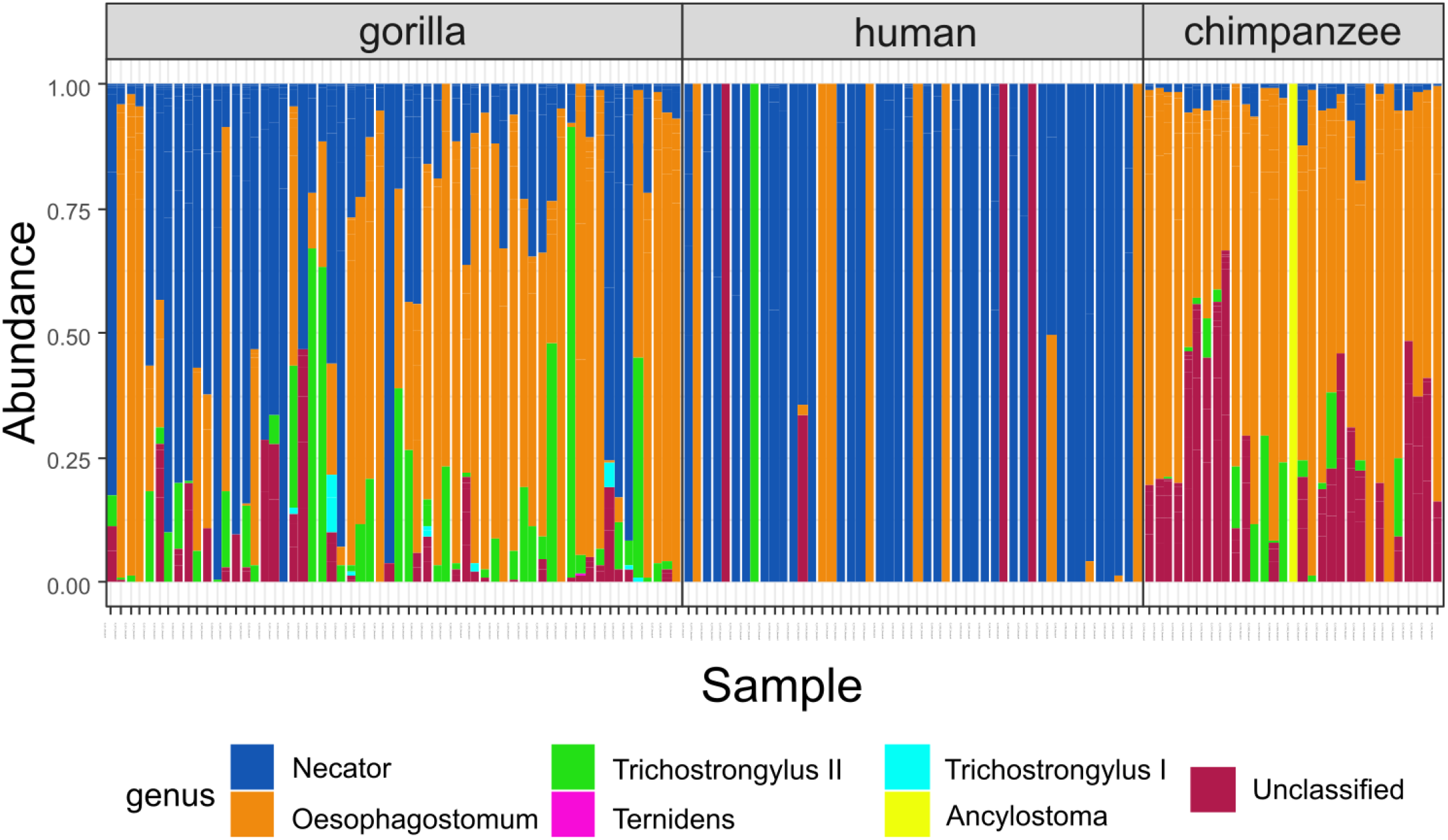
Bar graph showing relative community composition of strongylid nematodes in examined samples. Each column represents a sample. Relative abundances of reads are depicted as colour panels.

Variant diversity (*x′* = 7; min. = 1, max. = 17) differed among the studied hosts (GLM: F_(2,138)_ = 203.36, p < 0.0001). Variant diversity in humans was lower compared to both species of great apes (Tukey post-hoc testing: p = 0.0001 for all pairwise comparisons) **(Figure 2)**, while there was no evidence of significant differences between gorillas and chimpanzees (p > 0.3). PCoA diagrams based both on Jaccard and Bray-Curtis ecological distances confirmed clear differences between humans and great apes in both composition and relative abundance of strongylid ASVs **(Figure 3)**. Significant differences between different host species in the composition of their strongylid nematode communities were further confirmed by PERMANOVA (Jaccard: F_(2,138)_ = 11.655, p = 0.001; Bray-Curtis: F_(2,138)_ = 14.644, p = 0.001) and ANOSIM (Jaccard: R = 0.4456, p = 0.001; Bray-Curtis: R = 0.4204, p = 0.001) tests. Tukey post-hoc testing revealed significant differences between humans and other great apes both for Jaccard and Bray-Curtis (p < 0.01 for all pair-wise combinations) distances. Within great apes, there was no statistically significant result for Jaccard (p = 0.36) indicating roughly the same composition of strongylid ASVs; however, results for Bray-Curtis indicated differences in relative abundances (proportion) of ASVs between great apes (p < 0.001). Mvabund testing confirmed the interspecific differences (mvabund: ΔDF = 2, χ2 = 1002.371, p = 0.001) and identified 17 ITS-2 ASVs with whose different relative abundances were the main driving force of diversity between different host species **(Figure 4)**. Differences among hosts were mainly due to greater frequencies of *O. stephanostomum, N. gorillae, Trichostrongylus* type II and unclassified strongylids in great apes, whereas *N. americanus* was most frequent in humans.

**Figure 2:**
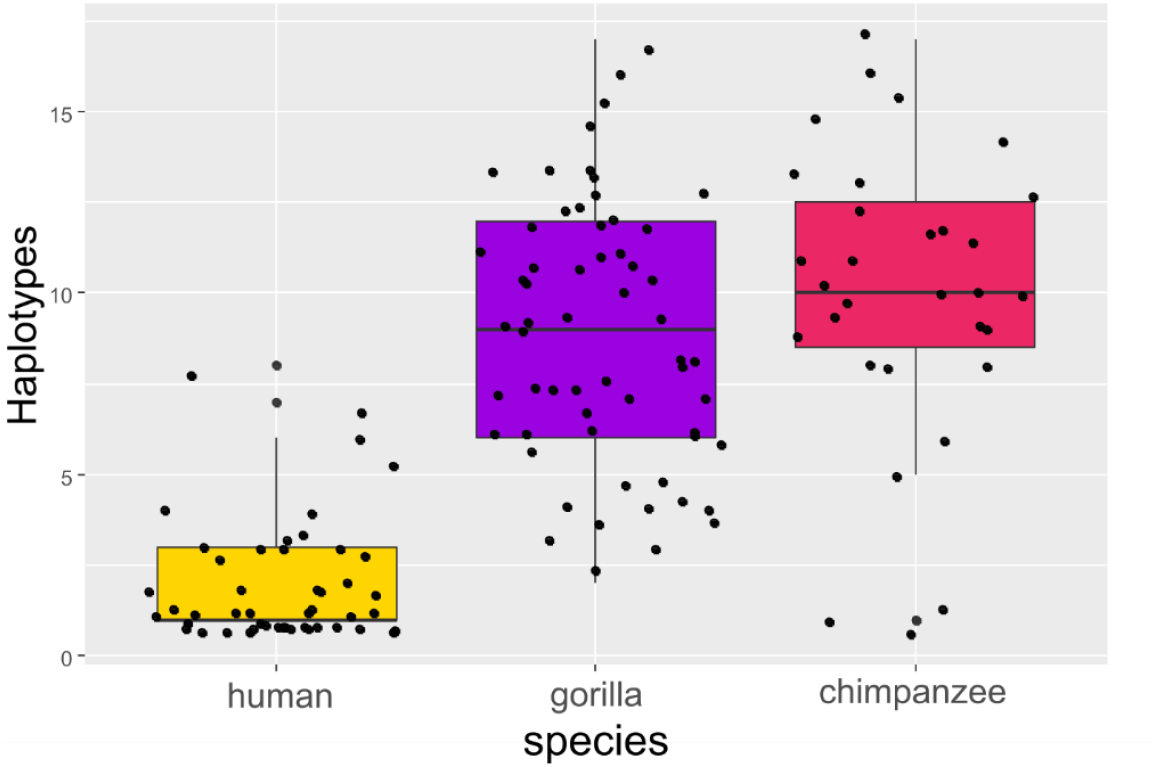
Alpha diversity of strongylid nematode communities, boxplot of amplicone sequencing variants’ (ASVs) counts for each sample (dots) according to host species. Different letters above boxes indicate statistically significant differences according to GLM test.

**Figure 3:**
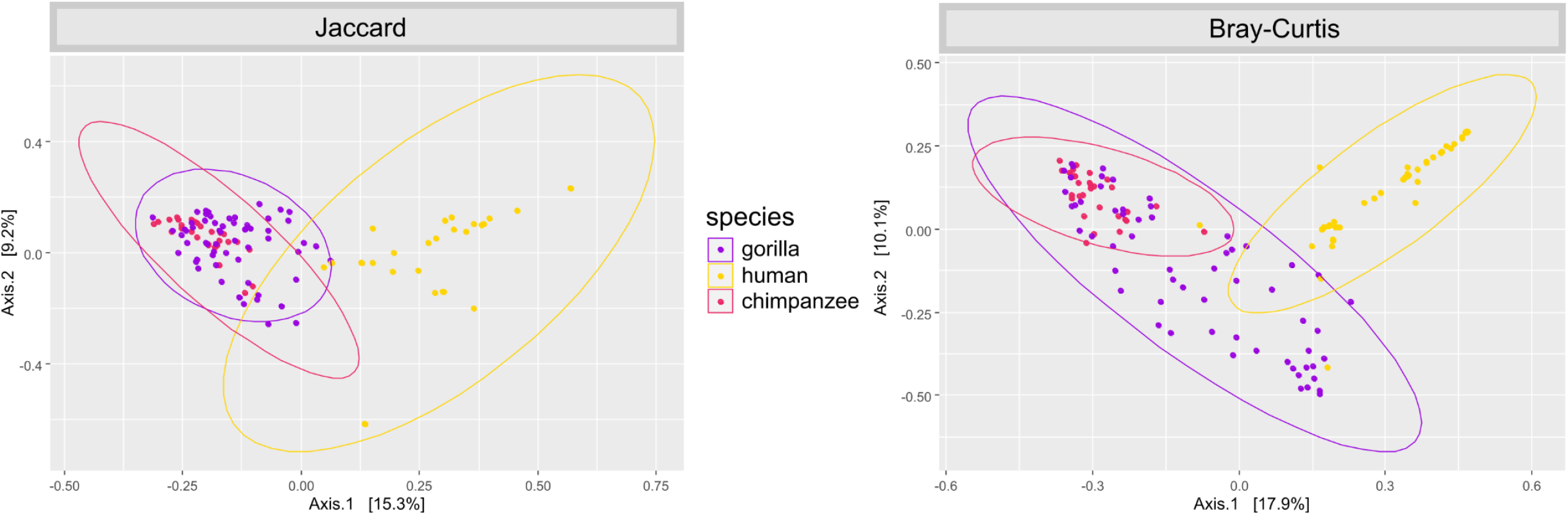
PCoA ordination diagrams of beta diversity of strongylid nematode communities based on Jaccard ecological distance: presence/absence of amplicone sequencing variants (ASVs); Bray-Curtis ecological distance (relative abundances of reads).

**Figure 4:**
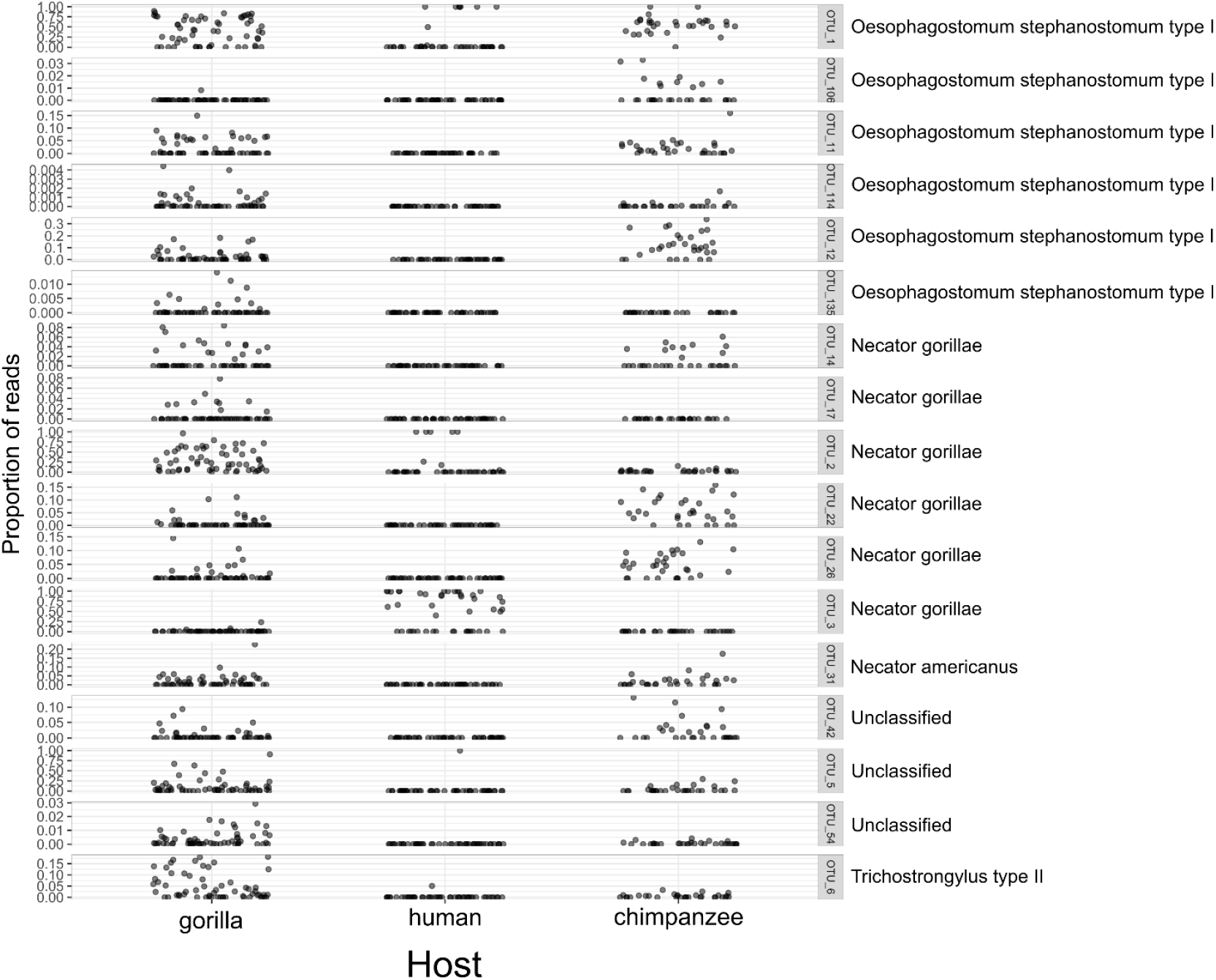
Plots showing relative abundance of ITS-2 amplicone sequencing variants (ASVs) indicated by Mvabund analyses as a driving force of differences among studied hosts.

We found no significant impact of behavioral or hygiene habits of the local people on either strongylid alpha or strongylid beta diversity (GLM: p > 0.05; PERMANOVA: p > 0.05; ANOSIM: p > 0.05).

## DISCUSSION

### Strongylid community composition

We explored the strongylid diversity and transmission patterns in humans and great apes sharing the same habitat in an unprotected area at the northern border of the Dja Faunal Reserve (Dja FR), Cameroon. Using the ITS-2 locus for identification, general taxonomic assignment revealed 95 strongylid ITS-2 amplicon sequence variants (ASVs), of which we could classify 65 at the genus/species level. In contrast to our previous study in Dzanga Sangha Protected Areas (DSPA), Central African Republic (CAR), where only two variants (from the total of 85) remained unassigned (Pafčo et al., 2019), our data from Dja contained 32 unassigned ASVs on the genus level; this suggests a more diverse strongylid fauna in Dja apes and humans, and further indicates that strongylid nematodes are a rather understudied group with unexplored diversity.

Overall, the composition of strongylid communities found in Dja remained generally consistent with previous studies, suggesting that *Necator* and *Oesophagostomum* are the most prevalent strongylid genera in African apes and humans (Cibot et al., 2015; Ghai et al., 2014; Hasegawa et al., 2017; Mason et al. 2022, Pafčo et al., 2018, 2019), but unlike in previous studies, these were followed by *Trichostrongylus* and unassigned genera. Dja apes were mostly infected by variants of *O. stephanostomum* and *N. gorillae*, both commonly found in great apes (Cibot et al., 2015; Ghai et al., 2014; Mason et al. 2022; Pafčo et al., 2019). Humans were mostly infected by *N. americanus* variants, confirming that *N. americanus* is the dominant human-specific hookworm in general (Hotez et al., 2005). Great apes exhibited higher strongylid diversity than humans and mixed infections of more than one strongylid species were frequently observed, which is consistent with previous findings in DSPA, CAR (Pafčo et al., 2019).

### Necator

We discovered six ASVs of *Necator* spp. (6.19 % from the total of 97 ASVs found) being shared between multiple hosts. Besides the human hookworm *N. americanus*, other *Necator* species (*N. exilidens, N. congolensis* and *N. gorillae*) have been reported in great apes (Cummins, 1912; Gedoelst, 1916; Noda & Yamada, 1964) and also in humans in Africa (Hasegawa et al., 2014; Kalousová et al., 2016; Pafčo et al., 2019). Four variants of *N. americanus* and two variants of *N. gorillae* were found co-infecting humans and great apes, suggesting ongoing transmission events previously described in the tropical forest ecosystem in DSPA, CAR (Hasegawa et al., 2014; Pafčo et al., 2018, 2019). While *N. americanus* variants were found mostly in humans, four of them were shared with gorillas; this demonstrates that *N. americanus* is not a solely human-specific parasite. Moreover, such a finding was previously observed in African great apes (Hasegawa et al., 2014; Mason et al. 2022; Pafčo et al., 2019). *Necator gorillae* variants were found predominantly in great apes, suggesting its probable ape origin, but it was also shared with humans. The *N. gorillae* variants corresponded to those previously found in gorillas in Gabon (Hasegawa et al., 2017). We did not find evidence of *N. americanus* infecting wild chimpanzees, which supports a previous hypothesis of a lower susceptibility of chimpanzees to *N. americanus* infections (Hasegawa et al., 2017), despite some cases of chimpanzee infections having been previously recorded (Orihel, 1971; Pafčo et al., 2019). Additionally, we found a few variants of undetermined *Necator* sp. in Dja apes corresponding to variant III-1 first found in humans in DPSA, CAR by Hasegawa et al. (2014), later reported in western lowland gorillas across several African localities (Hasegawa et al., 2017; Mason et al. 2022). Hasegawa et al. (2014) speculated that variant III-1 sequences may represent *N. congolensis* or *N. exilidens*, previously described in chimpanzees (Cummins, 1912; Gedoelst, 1916); however, the original descriptions of *N. congolensis* or *N. exilidens* were made at the beginning of the last century, and even “traditional” morphology-based taxonomy of *Necator* non-*americanus* species remain unclear (Kalousová et al., 2016). Several *Necator* species are clearly capable of infecting both humans and NHPs, at least in habitats where they share the same environment. However, the exact species diversity is not known, nor is the epidemiology and ability (particularly of the non-*americanus* species) to spread in human populations. Therefore, large-scale studies covering multiple populations of wild great apes, other NHPs and humans, with utilization of advanced HTS tools combined with modern morphological characterizations will be required for better understanding of *Necator* epidemiology.

### Oesophagostomum

Two *Oesophagostomum* species are commonly found in great apes and humans throughout Africa – *O. stephanostomum* (Cibot et al., 2015) and *O. bifurcum* in humans, especially in West Africa (Ziem et al., 2006); other *Oesophagostomum* species have been recorded, but they are much rarer (Modrý et al., 2018; Ota et al., 2015). We recorded one variant of *O. stephanostomum* type I shared among great apes and humans in Dja, corresponding to the variant infecting NHPs and humans in Kibale, Uganda (Cibot et al., 2015). This makes our finding the second observation of *O. stephanostomum* in humans providing evidence that *Oesophagostomum* species have zoonotic potential under suitable circumstances. Pafčo et al. (2019) also reported *O. stephanostomum* infecting NHPs in DSPA, CAR, but not in humans, thus suggesting its ape origin. This is also supported by Mason et al. (2022), who found *O. stephanostomum* in high prevalence in western lowland gorillas across several African localities. We only found the second *Oesophagostomum* group (*Oesophagostomum stephanostomum* type II) in great apes; this group was previously described in western lowland gorillas in Moukalaba-Doudau National Park (MDNP), Gabon (Makouloutou et al., 2014) and in Dja FR, Cameroon (using the same gorilla dataset as was used for this study; Mason et al., 2022). We only found the other variants of undetermined *Oesophagostomum* sp. in one gorilla. They correspond to *Oesophagostomum* sequences from humans and NHPs in Kibale, Uganda (Ghai et al., 2014), further recorded by Cibot et al. (2015) in olive baboons in other part of Kibale, Uganda. In contrast to Pafčo et al. (2019), who found *O. bifurcum* infecting NHP hosts in DSPA, CAR, we did not find evidence of this species in any studied host in Dja, although this species is known to be commonly infecting both humans and NHPs in Africa (Gasser, De Gruijter, & Polderman, 2006; Mason et al., 2022; Ziem et al., 2006).

### Other strongylids

Other strongylid nematodes may also infect humans and NHPs in Africa, such as the “false hookworm” *Ternidens deminutus*, the cyathostomine worm *Murshidia* spp. (Schindler, De Gruijter, Polderman, & Gasser, 2005), strongylids belonging to Trichostrongylidae (Durette-Desset, Chabaud, Ashford, Butynski, & Reid, 1992; Pafčo et al., 2019) and other pulmonary strongylids such as *Mammomonogamus* (Červená et al., 2017). We found several ASVs of *Trichostrongylus* spp. to be harbored by Dja great apes (their strongest BlastN matches were to trichostrongylids parasitic in sheep), and we found one variant to be shared between great apes and one Dja human, corresponding to the *Trichostrongylus* variant from chimpanzees living in degraded forest fragments in Bulindi, Uganda (McLennan, Hasegawa, Bardi, & Huffman, 2017). Variants of *Trichostrongylus* were reported by Pafčo et al. (2019) and Mason et al. (2022) in lowland gorillas, and adult *Trichostrongylus* worms were found in necropsied mountain gorillas in Rwanda (Hastings, 1992). Although several cases of *Trichostrongylus* infections have been reported in humans in north-eastern Thailand, Lao People’s Democratic Republic (PDR) and urban areas of Salvador City, Brazil (Phosuk et al., 2013; Souza et al., 2013), human *Trichostrongylus* infections are considered rather incidental. We found one variant of *Ternidens deminutus* infecting western lowland gorillas, closely similar to the one from Mona monkeys (*Cercopithecus mona*) found in Ghana (Schindler et al., 2005); this finding also corresponded to Pafčo et al. (2019), who found four *T. deminutus* variants infecting great apes, being closely related to the same sequence. *Ternidens deminutus* is considered to be a neglected parasite of humans (Schindler et al., 2005) and has also been reported in chimpanzees of Tai, Côte d’ Ivoire (Metzger, 2014) and in western lowland gorillas of Loango National Park, Gabon and in DSPA, CAR by Mason et al. (2022) raising questions about its origin and zoonotic potential. *Ancylostoma* is considered a human-specific parasite and was found by Pafčo et al. (2019) in humans in DSPA, Central African Republic. Our data show evidence for the first chimpanzee infection by *Ancylostoma* sp. ever recorded; however, we could not specifically assign the variant to a known *Ancylostoma* species, and it was found only in one chimpanzee sample, representing 100% of total sample reads. Such homogeneity in chimpanzees is rather unusual, according to our dataset.

### Zoonotic patterns

In Dja, humans exhibited lower strongylid alpha diversity than great apes and formed a separate cluster distinct from great apes, which was caused by dominance of *N. americanus* variants in both prevalence and relative abundance (measured as the proportion of sequencing reads assigned to this species). On the other hand, the strongylid communities of the two great apes species overlapped and were dominated by variants belonging to *N. gorillae, O. stephanostomum, Trichostrongylus* type II and unclassified variants. Our results corroborated the results from DSPA, CAR (Pafčo et al., 2019), where the composition of strongylid communities was also shaped by the extent of habitat sharing, which is much more intense among species of great apes than between humans and great apes. Infective larvae (L3) of monoxenous strongylid nematodes develop in the external environment (Anderson, 2000), thus habitat sharing increases the risk of infection and transmission between the hosts. In both localities, the observed pattern of strongylid communities did not reflect the phylogenetic relationships of the hosts and rather reflected ecosystem sharing rate as the community is more similar between great apes than between humans and chimpanzees. On the contrary, Vlčková et al. (2018) observed a higher alpha diversity of *Entamoeba* (protozoan parasite) communities in humans compared to great apes in the same locality. Furthermore, interestingly, the composition of human and chimpanzee *Entamoeba* communities overlapped, while that of gorillas formed a clearly separated cluster, displaying a pattern that reflects the phylogenetic distance between the hosts. Mann et al. (2020) analysed gut protists and nematodes of NHPs from various sites using the 16S phylogenetic marker. Although 16S markers cannot provide high phylogenetic resolution for strongylid nematodes (Aivelo & Medlar, 2017), these results also showed that gut eukaryotes (unlike symbiotic gut microbes) were only weakly structured by primate phylogeny, although other parts of the gut microbiome (such as mycobiome) can be more plastic and can reflect host phylogeny (Sharma et al., 2022).

We recorded higher numbers of strongylid ASVs shared between humans and great apes in Dja FR in comparison to DSPA, CAR. The majority of human respondents in Dja followed a more agriculture-oriented lifestyle (Table 3: 54.17 % stated “farmer” as their occupation) while in DSPA, CAR the studied humans were hunter-gatherers (Pafčo et al., 2019). Moreover, Dja FR is under high anthropogenic pressure. The forest is degraded, and fragmented, as intense logging, hunting and crops occur in the area (Arlet & Molleman, 2010; Ávila et al., 2019; Betti, 2004; Tagg et al., 2018). Based on previous studies from different localities, we assume that the agricultural fields in Dja can attract animals (such as chimpanzees), resulting in crop-raiding with defecation (Arlet & Molleman, 2010; Tudge, Brittain, Kentatchime, Kamogne Tagne, & Rowcliffe, 2022). Local people often walk barefoot through Dja agricultural fields and eat crops straight from the ground without previously washing them **(Table 3)**. Together with almost no anthelmintic treatment and poor sanitation, the transmission of strongylid parasites can be greatly facilitated as *Necator, Oesophagostomum* and *Trichostrongylus* are parasites transmitted by skin penetration or oral ingestion (Anderson, 2000). Sub-Saharan African humans have always shared their habitat with NHPs, but the situation has changed dramatically in recent decades, as the human population is growing rapidly, and their requirements for space and resources are much higher than before. Our results here indicate that increased spatial proximity of wildlife and humans creates an opportunity for pathogen spillover in both directions.

**Table 3:** Results of questionnaires based on respondents answers. Survey focused mainly on human-animal interactions, lifestyle and hygiene standards.

| Activity | Yes | No | Frequently | Sometimes | Never | River | Well | Both |
| --- | --- | --- | --- | --- | --- | --- | --- | --- |
| **Entering the forest** | - | - | 64.6 % | 29.2 % | 6.2 % | - | - | - |
| **Contact with wild apes** | 89.6 % | 10.4 % | - | - | - | - | - | - |
| **Contact with apes' feces** | 70.8 % | 29.2 % | - | - | - | - | - | - |
| **Wild apes around household** | 22.9 % | 77.1 % | - | - | - | - | - | - |
| **Wearing shoes in the forest** | 39.6 % | 60.4 % | - | - | - | - | - | - |
| **Eating from the ground** | 97.9 % | 2.1 % | - | - | - | - | - | - |
| **Washing crops before eating** | 8.3 % | 91.7 % | - | - | - | - | - | - |
| **Washing hands before eating** | 37.5 % | 62.5 % | - | - | - | - | - | - |
| **Drinking water sources** | - | - | - | - | - | 89.6 % | 2.1 % | 8.3 % |
| **Anthelmintic treatment** | 29.2 % | 70.8 % | - | - | - | - | - | - |
| **Taking plant-based drugs** | 4.2 % | 91.7 % | - | - | - | - | - | 4.1 % |

On the other hand, Mason et al. (2020) observed lower strongylid diversity in western lowland gorillas in Dja FR compared to other study areas (including DSPA, CAR), which may indicate the impact of greater anthropogenic disturbance in Dja FR on strongylid communities. Traditionally, parasites are thought to have negative effects on the host; however, they are a natural part of the host environment due to millions of years of evolution (Perry, 2014). It appears that a loss of parasitic symbionts occurs in areas of increased anthropogenic pressure (Barelli et al., 2020). It is speculated that the loss of parasitic symbionts in humans may contribute to an increase in autoimmune diseases (Maizels, 2016). We have demonstrated that parasite transmission is likely to be higher in unprotected areas such as in Dja than in protected areas such as in Dzanga-Sangha National Park (DSPA, CAR), but we must keep in mind that increased anthropogenic pressure may also alter or even reduce parasite diversity, with consequences that are as yet incomprehensible to us.

## CONCLUSION

We revealed complex strongylid nematode communities of great apes and humans sharing an unprotected tropical forest habitat in Cameroon. The great apes exhibited a greater diversity of the strongylid fauna harbouring more amplicon sequencing variants (ASVs) and rare variants in comparison to humans. *Oesophagostomum* and *Necator* were the dominant components of strongylid communities in all studied hosts, and the driving force of strongylid overlaps. Human communities were dominated by *Necator americanus*; although generally thought to be human-specific, this parasite was also shared by gorillas. *Necator gorillae*, originally thought to be a parasite confined to NHPs, was widespread across all studied host species, including humans. We observed a second case of *O. stephanostomum* infection in humans. In contrast to previous studies conducted in the DSPA, CAR, we recorded more genera and variants being shared between humans and great apes, probably due to significant anthropogenic pressure in the periphery of the reserve, which is not protected. Most African apes occur outside protected areas (Carvalho et al., 2021) and thus improving the effectiveness of pathogen monitoring, conservation efforts and management not only inside, but also outside, protected areas is urgently warranted.

## ACKNOWLEDGEMENTS

This study was supported by the Czech Science Foundation (18-24345S and partially also by 22-16475S); Masaryk University (MUNI/A/1488/2021) and Institute of Vertebrate Biology, Czech Academy of Sciences (RVO:68081766). Further support for work by V.I. was provided by the Fulbright Foundation (fellowship number PS00299111). We would like to express our gratitude also to the Ministère de la Recherche Scientifique et de l’Innovation a Ministère des Forêts et de la Faune, Cameroon for permission to conduct the research in Cameroon; Antwerp Zoo Society, Belgium and Project Grands Singes, Cameroon for welcoming the project and logistical support in the field; and all local trackers and assistants and other people, who helped with sample collection: Arlette Tchankugni Nguemfo, Klára Vlčková, Zuzana Tehlárová, Dagmar Jirsová. We acknowledge the CF Genomics CEITEC MU supported by the NCMG research infrastructure (LM2015091 funded by MEYS CR) for their support with obtaining scientific data presented in this paper. We would like to acknowledge Core Facility - Genomics within CEITEC, Masaryk University for technical support and valuable advice. Computational resources were supplied by the project “e-Infrastruktura CZ” (e-INFRA CZ LM2018140) supported by the Ministry of Education, Youth and Sports of the Czech Republic.

## AUTHOR CONTRIBUTIONS

B.P. and V.I. designed the study, carried out the molecular laboratory work and drafted the manuscript. B.P., Do.M., Charmance and N.T. carried out/secured sample and data collection. V.I. performed the bionformatic and statistical analysis under B.P and J.K.’s supervision. E.M.S. and K.J.P. significantly improved the manuscript. N.T. made English corrections. K.J.P. and D.M. secured funding for the study, with some support for V.I. via E.M.S. All authors approved final publication.

## DATA ACCESSIBILITY AND BENEFIT-SHARING

Sequencing data are available under accession number of the whole project PRJEB34054. Accession numbers related to BlastN hits for our sequences are listed in Table 2. Research involved many scientists around the world and collaborators from surveyed localities, added as co-authors to this manuscript. The results were shared with all collaborators and will be useful for broad scientific community, as mentioned above.

